# Comprehensive Analysis of Indels in Whole-genome Microsatellite Regions and Microsatellite Instability across 21 Cancer Types

**DOI:** 10.1101/406975

**Authors:** Akihiro Fujimoto, Masashi Fujita, Takanori Hasegawa, Jing Hao Wong, Kazuhiro Maejima, Aya Oku-Sasaki, Kaoru Nakano, Yuichi Shiraishi, Satoru Miyano, Seiya Imoto, Michael R Stratton, Steven G Rosen, Hidewaki Nakagawa, ICGC/TCGA Pan-Cancer Analysis of Whole Genomes Network

**Affiliations:** Laboratory for Cancer Genomics, RIKEN Center for Integrative Medical Science, Japan; Department of Drug Discovery Medicine, Kyoto University Graduate School of Medicine, Japan; Health Intelligence Center, Institute of Medical Sciences, The University of Tokyo, Japan; Human Genome Center, Institute of Medical Sciences, The University of Tokyo, Japan; Welcome Trust Sanger Institute, UK; Center for Computational Biology, Duke-NUS Medical School, Singapore

## Abstract

Microsatellites are repeats of 1-6bp units and ∼10 million microsatellites have been identified across the human genome. Microsatellites are vulnerable to DNA mismatch errors, and have thus been used to detect cancers with mismatch repair deficiency. To reveal the mutational landscape of the microsatellite repeat regions at the genome level, we analyzed approximately 20.1 billion microsatellites in 2,717 whole genomes of pan-cancer samples across 21 tissue types. Firstly, we developed a new insertion and deletion caller (MIMcall) that takes into consideration the error patterns of different types of microsatellites. Among the 2,717 pan-cancer samples, our analysis identified 31 samples, including colorectal, uterus, and stomach cancers, with higher microsatellite mutation rate (≥ 0.03), which we defined as microsatellite instability (MSI) cancers in genome-wide level. Next, we found 20 highly-mutated microsatellites that can be used to detect MSI cancers with high sensitivity. Third, we found that replication timing and DNA shape were significantly associated with mutation rates of the microsatellites. Analysis of germline variation of the microsatellites suggested that the amount of germline variations and somatic mutation rates were correlated. Lastly, analysis of mutations in mismatch repair genes showed that somatic SNVs and short indels had larger functional impact than germline mutations and structural variations. Our analysis provides a comprehensive picture of mutations in the microsatellite regions, and reveals possible causes of mutations, as well as provides a useful marker set for MSI detection.

## Introduction

Recent large-scale whole genome sequencing studies have revealed the complexity of the mutational landscape of the cancer genome (1-4). In cancer genomes, various types of mutations, such as SNVs (single nucleotide variants), short indels (insertions and deletions), genomic rearrangements, copy number alterations, insertion of retrotransposons, and virus genome integrations, have been identified, and their oncogenic roles have been characterized (1-5). Additionally, genome sequencing studies have revealed the molecular basis of somatic mutations (6-9). However, somatic mutations in microsatellites or repeat sequences have not been well-characterized in a large whole genome sequencing cohort due to difficulties in accurately detecting mutations using presently available short-read sequencing technologies.

A microsatellite is defined as a tract of repetitive DNA motif composed of short repeating units (10). The mutation rate of microsatellites has been known to be higher than other genomic regions due to DNA polymerase slippage during DNA replication and repair (10). Due to their fragility, microsatellites are used as markers of genomic instability in cancer (11). In cancer genetics studies, microsatellite instability (MSI) has been used for molecular diagnosis of Lynch syndrome and cancers with mismatch repair deficiency (11). Furthermore, MSI-positive tumors are generally burdened with higher numbers of somatic mutations and present many mutation-associated neo-antigens, which might be recognized by the immune system. Presently, MSI can also be used as a marker to predict the effect of immune therapy (12). The MSI phenotype is most common in colorectal cancers, stomach cancers and uterine endometrial cancers (10-15%), although it has also been observed across many tumor types at a few % (11). The MSI phenotype is defined by the presence of somatic indels of the 2-5 microsatellite makers, whereby BAT25/26 mononucleotide microsatellites are widely used to establish MSI status (11).

Irrespective of the clinical importance of microsatellite, large-scale analysis of somatic changes in microsatellites across various type of cancers is limited for whole genome sequencing (WGS) data (13, 14). In the current study, we analyzed indels in microsatellites for 2,913 ICGC pan-cancer samples from 21 tissues (15) to reveal the whole genome mutational landscape of microsatellite regions. We developed a method to detect somatic indels in microsatellite regions, selected an appropriate parameter for our purpose, and identified indels in microsatellite regions. We identified MSI-positive samples and factors affecting the mutation rate of microsatellites, as well as highly-mutated microsatellites. We also analyzed the association of mutation rate of microsatellites with somatic and germline mutations in DNA repair genes, and compared mutational signatures between MSI and other samples. Our analysis provides a comprehensive picture of mutations in the microsatellite regions, and reveals possible causes of mutations, as well as provides a useful marker set for MSI detection.

## Results

### Identification of microsatellite regions in the genome

We detected microsatellites using three methods (MsDetector, Tandem Repeat Finder, and MISA software) (16-18). To exclude microsatellites potentially arising from read mapping errors, we selected microsatellites based on the uniqueness of flanking sequences and pattern of repeats. A total of 9,292,677 microsatellites were used for subsequent analyses. Within these selected microsatellites, it was observed that the MISA software identified a larger number compared to other methods (**Supplementary Fig. 1**).

**Figure 1.**
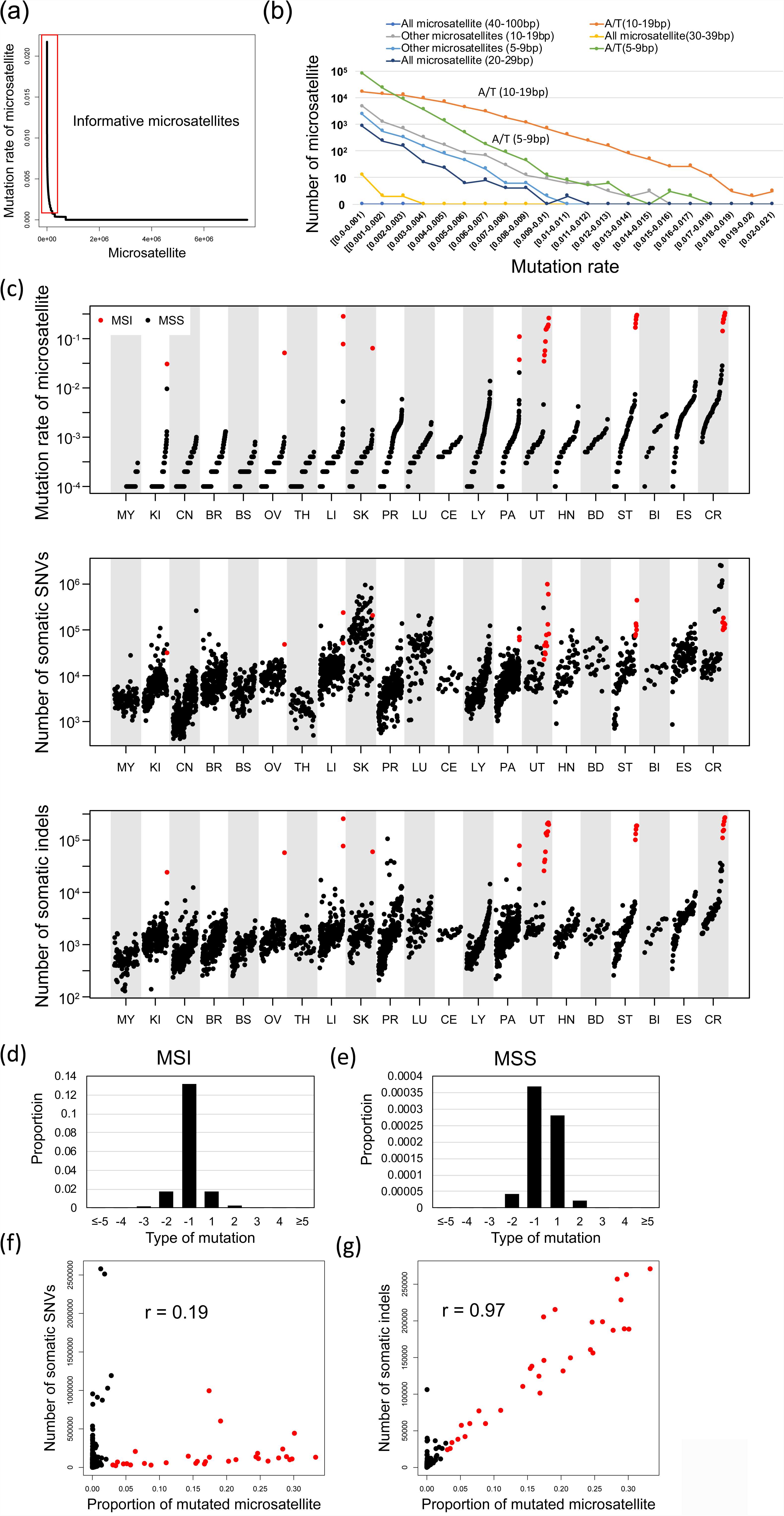
Pattern of somatic indels in microsatellite regions. **(a)** Mutation rate of each microsatellite. 7,650,128 microsatellites were sorted by the proportion of mutated samples. The red box indicates informative microsatellites defined in this study (proportion of mutated samples ≥ 0.001). **(b)** Mutation rate among different microsatellites. **(c)** Comparison of mutation rate of microsatellites, number of somatic SNVs and indels in different types of cancer. MSI samples are shown in red. **(d-e)** Pattern of insertions (positive change in repeat length in x-axis) and deletions (negative change in repeat length in x-axis) in microsatellites between the MSI **(d)** and MSS samples **(e)**. Correlation between the mutation rate of microsatellites and the number of somatic SNVs. Pearson’s product-moment correlation; *r* = 0.19, *p-value* = 1.6×10^-23^. MSI samples are shown in red. **(f)** Correlation between the mutation rate of microsatellites and the number of somatic indels. Pearson’s product-moment correlation; *r* = 0.97, *p-value* < 1.0×10^-200^. MSI samples are shown in red.

### Error rate estimation of microsatellites

During library preparation and sequencing processes, indel errors can be introduced by PCR in reads containing short repeats, due to replication slippage of DNA polymerases. Since the error rates should depend on the length and type of microsatellites, we first estimated the error rates of different types of microsatellites. The type of microsatellites was defined by length of the microsatellite region in the reference genome and repeat unit (see **Materials and Methods**). Using sequence data of chrX from 32 normal tissues of male individuals, we estimated the error rate among different types and lengths of microsatellites. As the male chrX is hemizygotic, the error rate can be inferred without the influence of heterozygous polymorphisms (19, 20). As expected, error rates depended on the unit and length of the microsatellites, with longer microsatellites having higher error rates (**Supplementary Fig. 2**). In all types of microsatellites, deletion errors were more frequent than insertion errors, and smaller changes of unit number were predominant (**Supplementary Fig. 2**). These results suggest that PCR or sequencing processes are prone to induce short deletion errors. Error rates of microsatellites between 5-9bp in length within the reference genome were very low (< 0.2%), while those of longer microsatellites were higher (> 5% error rate for 20-100bp of microsatellites in the reference genome length). Two bp repeats had higher error rates than other microsatellites (**Supplementary Fig. 2**). The A/T type of microsatellite was observed to have higher error rates compared to G/C microsatellites (**Supplementary Fig. 2**). Since the estimated error rates were quite different among the types and lengths of microsatellites, we used the difference between error rates to detect somatic indels in the microsatellites. We generated a table of error rates for analyzing mutations in microsatellite regions based on the estimated error rates (**Supplementary Table 2**).

**Figure 2.**
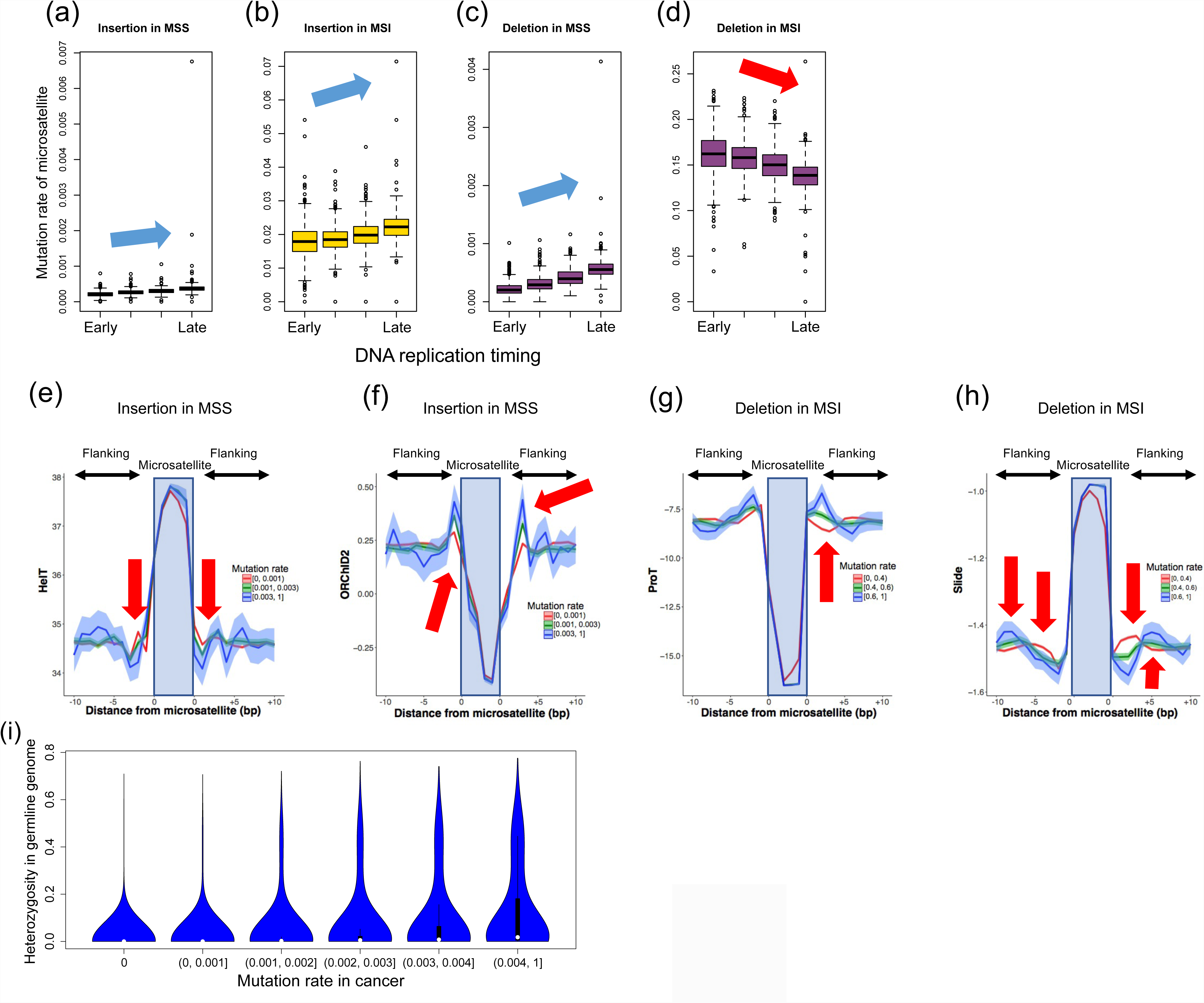
Analysis of mutation rate of each microsatellite. The mutation rate of 198,578 informative microsatellites was analyzed. **(a-d)** Association of replication timing with insertion and deletion rates. The edges of the boxes represent the 25th and 75th percentile values. The whiskers represent the most extreme data points, which are no more than 1.5 times the interquartile range from the boxes. **(a)** Insertions in MSS samples (𝒳^2^ test; *p-value* < 1×10^-200^), **(b)** Insertions in MSI samples (𝒳^2^ test; *p-value* < 1×10^-200^), **(c)** Deletions in MSS samples (𝒳^2^ test; *p-value* < 1×10^-200^), **(d)** Deletions in MSI samples (𝒳^2^ test; *p-value* < 1×10^-200^). **(e-h)** Association of DNA shape with mutation rate. Top 1000 A/T microsatellites with 10-30bp length were used for the analysis. The microsatellites were divided into the three categories based on the mutation rate. **(e)** HelT (Helix twist) of insertions in MSS samples. **(f)** ORChID2 (the ·OH Radical Cleavage Intensity) of insertions in MSS samples, **(g)** ProT (Propeller Twist) of deletions in MSI samples, **(h)** Slide of deletions in MSI samples. In this figure, we divided the microsatellites with mutation rates (0-0.001, 0.001-0.003, and > 0.003 for MSS, and 0-0.4, 0.4-0.8, and > 0.8 for MSI), and showed the DNA shape values. The arrows show base positions with significant association between the DNA shape values and mutation rates (**Supplementary Table 5**). **(i)** Germline variation and somatic mutation rates of microsatellites. The amount of germline variation was estimated using heterozygosity of normal tissues. X-axis; the proportion of mutated samples in each microsatellite (somatic mutation rate). Y-axis; heterozygosity of each microsatellite. Both rates were positively correlated (Pearson’s product-moment correlation; *r* = 0.31, *p-value* < 1.0×10^-100^).

### Validation with simulation data sets and setting of thresholds

Mutations in microsatellite regions were identified based on likelihoods (see Materials and Methods). To estimate false positive and false negative rates, and to select appropriate parameters, we generated simulation data sets by using sequence reads mapped on chrX of male individuals. First, we determined the genotype of each microsatellite on chrX. Since chrX is hemizygotic in male, we considered the dominant reads as the true genotype of each sample. We then mixed chrX reads from two male individuals and identified variations in the microsatellite regions with our algorithm. By comparing the true genotypes and genotypes from the mixed data, we estimated the false positive and negative rates. The false positive and negative rates were varied according to the likelihood values (*L*), and higher *L* had higher false negative and lower false positive rates (**Supplementary Fig. 3a, b, c**). To identify somatic mutations in microsatellites, we require reads that completely cover target microsatellites. The length of reads is about 100bp, therefore, longer microsatellites have fewer reads covering them compared to shorter microsatellites, and thus have a lower sensitivity (**Supplementary Fig. 3d**). Based on the analysis, *L* was set to −8 for cancer samples and −1 for matched normal samples.

**Figure 3.**
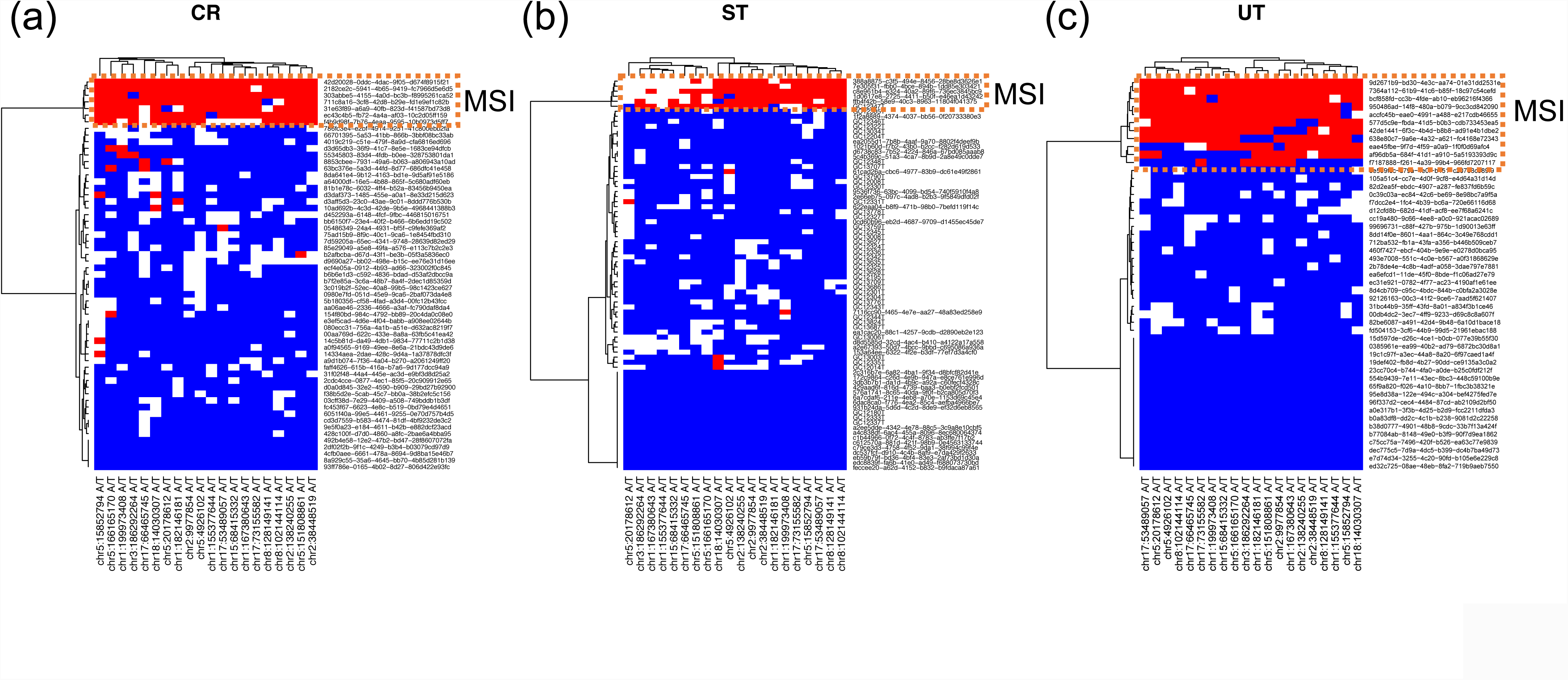
Highly-mutated microsatellite markers. Result of clustering analysis with the top 20 microsatellites. **(a)** CR (Colon/Rectum) cancer, **(b)** ST (Stomach) cancer, **(c)** UT (Uterus) cancer. **(d)** Result of validation study in the independent colon cancer cohort (n=36). MSI was defined by BAT25 or BAT26 (upper panel). Twelve new microsatellite markers (middle panel) were successfully amplified and analyzed for indel mutations by MiSeq (**Supplementary Fig. 12**). Fourteen highly-mutated microsatellites or homopolymers in coding regions were analyzed (lower panel). These mutations occurred specifically in MSI colorectal cancers. Mutation status are shown in brown (mutated) and white (WT; wild type) in the panel. Number of mutated samples in the MSI samples are shown in bar plot (right).

### Analysis of indels in the microsatellite regions in pan-cancer samples

We analyzed the whole genome sequence data of 2,917 pan-cancer samples (21) with our method and compared them against somatic and germline variants detected by the Pan-Cancer Analysis of Whole Genomes (PCAWG) project (22). To compare our results with somatic consensus indels from the four PCAWG indel callings, we gathered indels located ±5bp in the microsatellite regions in the PCAWG calls. On average, 1826.5 indels were detected by our indel caller (MIMcall) in the microsatellite regions. Of these, 1185.1 were found only by MIMcall (**Supplementary Fig. 4a**), suggesting a higher sensitivity of our indel calls compared to the other PCAWG callers for microsatellite regions. PCAWG calls removed repetitive regions to achieve highly accurate mutation calling, therefore our result can complement the PCAWG calls. We then compared the number of indels in the microsatellite regions between our indel caller and PCAWG callers. In the microsatellite regions, the number of indels uniquely identified by our indel caller was significantly correlated with that of commonly identified indels (identified by ≥ two PCAWG callers) (Pearson’s product moment correlation coefficient; *r* = 0.90, *p-value* < 10^-16^) (**Supplementary Fig. 4b**). We further performed experimental validation with Japanese liver cancer samples for the mutation candidates in longer microsatellites by capillary electrophoresis (**Supplementary Fig. 5**). The false discovery rate of our method was estimated to be 7% (2/29) (**Supplementary Fig. 5**). These results indicate that MIMcall can identify indels in the microsatellite region effectively.

**Figure 4.**
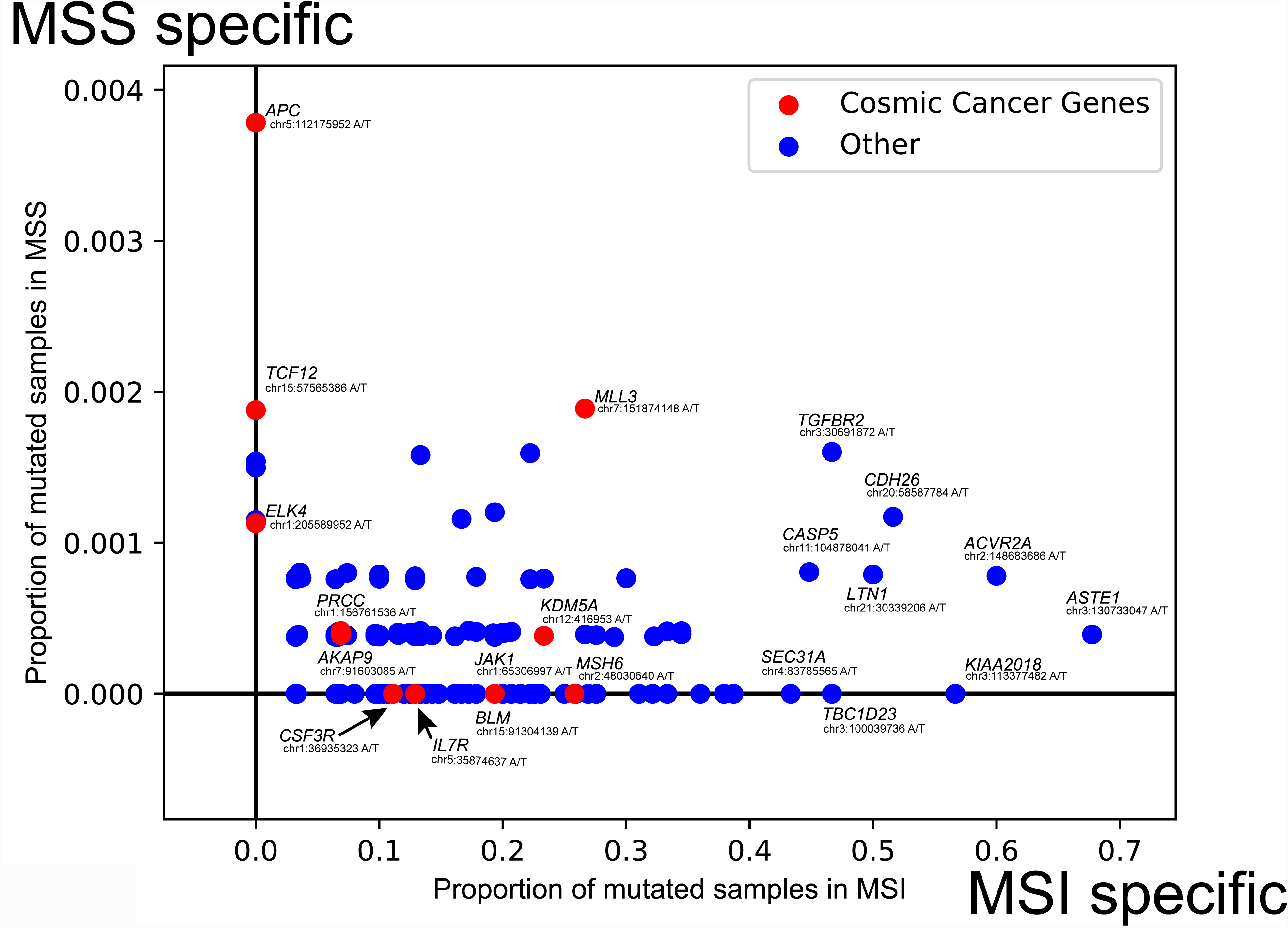
Proportion of mutated samples in coding microsatellites. x-axis; proportion of mutated samples in the MSI samples. y-axis; proportion of mutated samples in the MSS samples. Cosmic cancer genes are shown in red.

**Figure 5.**
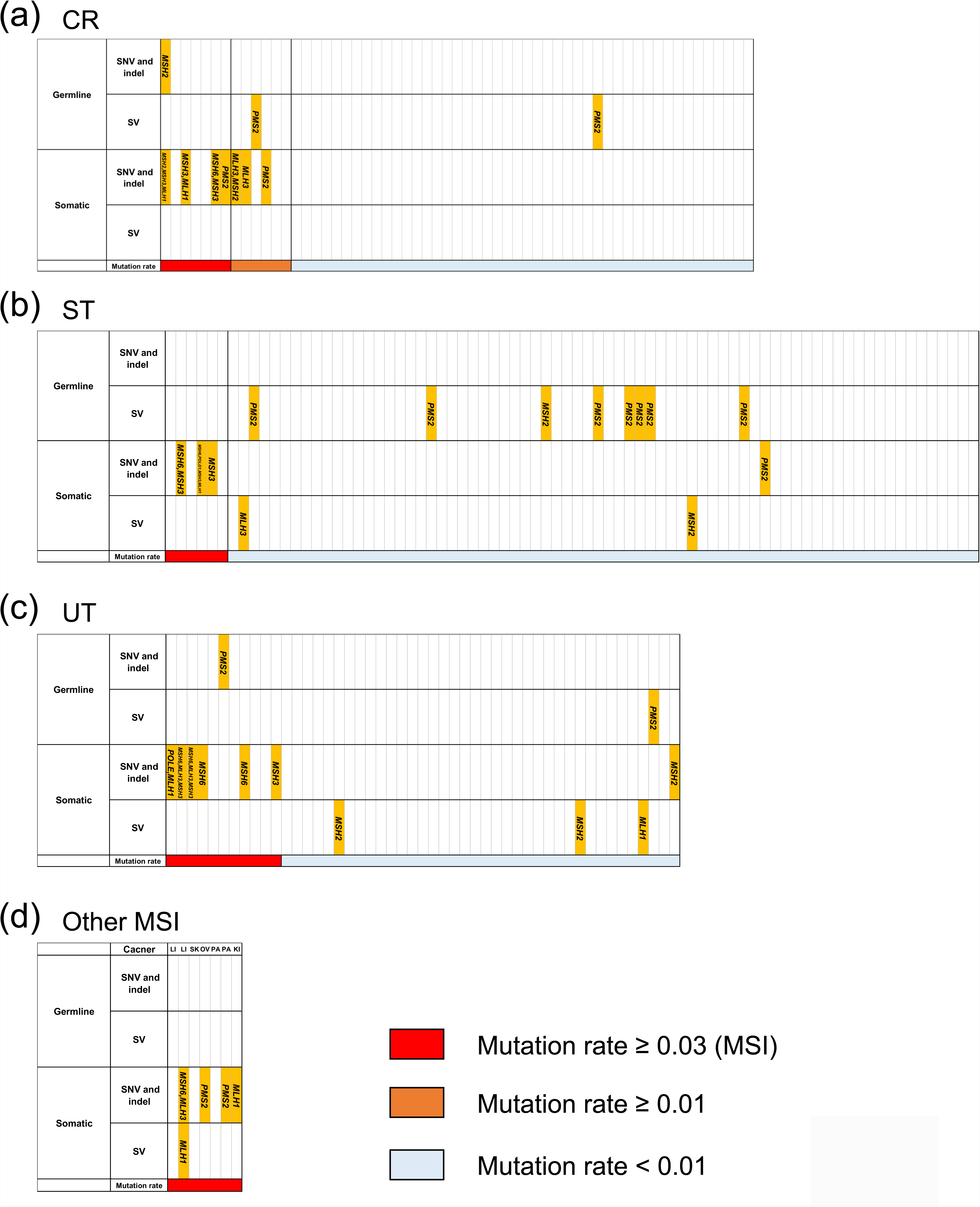
Mutation in mismatch repair (MMR) and proof-reading genes. Mutated genes, and mutation rate of microsatellites are shown.

Microsatellites covered by ≥ 15 reads in ≥ 2,500 samples (7,650,128 microsatellites) were subjected to further analysis, and samples with ≥ 6,000,000 testable microsatellites were used for the analysis (2,717 samples) (**Supplementary Table 3**). On average, 7,407,000 microsatellites were analyzed in each sample. We compared the proportion of mutated samples for each microsatellite. Most of the microsatellites in whole genomes were not mutated in the pan-cancer samples (**Fig. 1a**), we therefore selected 198,578 microsatellites with proportions of mutated samples ≥ 0.001 (more than 2-3 mutated samples in the pan-cancer samples) and considered them as informative microsatellites. The proportions of mutated samples were different among the types and lengths of microsatellite, with A/T microsatellites more frequently mutated than other types (**Fig. 1b**). The proportions of mutated microsatellites were significantly lower in exonic and intronic regions, but higher in non-genic regions (**Supplementary Fig. 6**). Microsatellites in CDS (coding sequence) regions would evolve to be more stable to avoid mutations and this would cause lower mutation rates in the CDS regions. Lower mutation rates in intronic regions suggests the influence of transcription-coupled repair.

**Figure 6.**
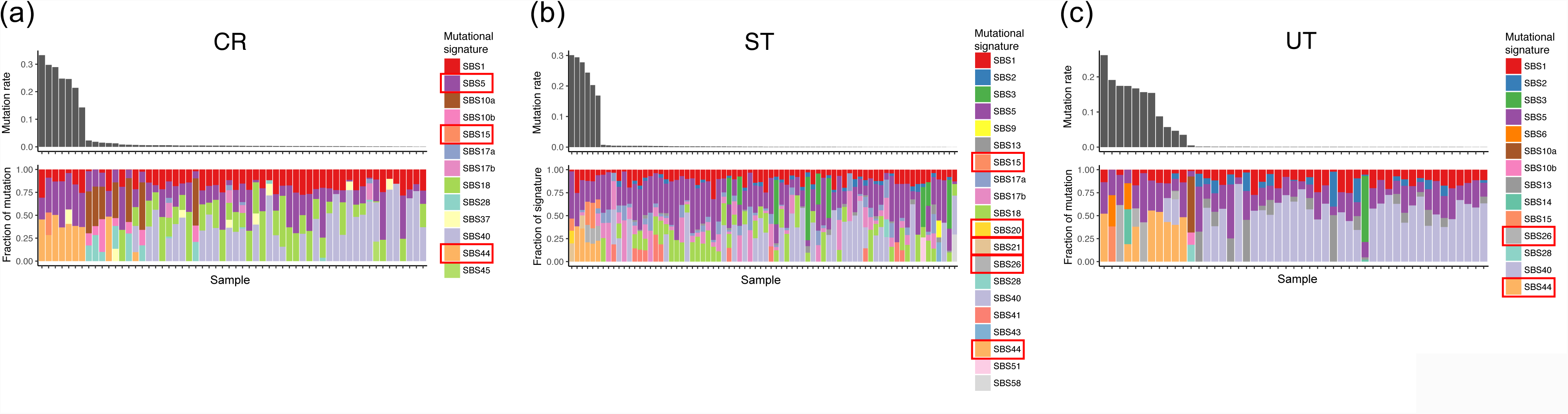
Comparison of mutational signatures between the MSI and MSS samples. **(a)** CR (Colon/Rectum) cancer, **(b)** ST (Stomach) cancer, **(c)** UT (Uterus) cancer. Signatures showing significant difference between the MSI and MSS samples are shown in rectangles in the legends (Wilcoxon signed-rank test, *q-value* < 0.05).

We next defined samples with proportions of mutated microsatellites ≥ 0.03 as microsatellite instability (MSI) samples (n=31) in genome-wide level (**Fig. 1c**), and others as microsatellite stable (MSS) samples. As expected, colorectal (CR), uterus (UT), and stomach (ST) cancers had a larger number of MSI samples, but MSI was also observed in a minority of samples for liver (LI), pancreas (PA), ovary (OV), kidney (KI), and skin (SK) cancers (**Fig. 1c**). The proportions of MSI samples were 11.9% for colorectal (7/59, 95% C.I. 4.9%-22.9%), 7.7% for stomach (6/78, 95% C.I. 2.8%-16.0%) and 22.0% for uterine (11/50, 95% C.I. 11.5%-36.0%) cancers, which are consistent with previous studies (23-25). We additionally analyzed Bethesda markers, which is a conventional marker set for MSI definition. However, the number of sequence reads that mapped to these regions was quite small and we could not analyze their mutation in most of the WGS (**Supplementary Fig. 7**).

The mutation pattern of microsatellites was different between the MSI and MSS samples. In the MSI samples, deletions were more predominant compared to insertions (**Fig. 1d and 1e**). We compared the proportion of mutated microsatellites, the number of somatic SNVs, and somatic indels (**Fig. 1f and 1g**). Although the numbers of SNVs were not strongly correlated with the proportion of mutated microsatellites (*r* = 0.19), the number of somatic indels were clearly correlated (*r* = 0.97) (**Fig. 1f and 1g**), suggesting that microsatellite and non-microsatellite indels are affected by common mechanisms of mutation and repair, and that the analysis of microsatellites can predict samples with large number of non-microsatellite indels.

### Mutability of microsatellite

Recent studies have suggested that epigenetic factors, such as DNA structures, and sequence motif, influence the mutation rate (8, 9, 26, 27). However, little is known about factors that influence the mutability of microsatellites. We first analyzed the replication timings and microsatellite mutation rates (proportion of mutated samples for a microsatellite) (**Fig. 2a-d**). The late-replicating regions had lower mutation rates for insertions and deletions in MSS samples and insertions in MSI samples (**Fig. 2abc**). However, an inverse pattern was observed for deletions of the MSI samples; the early replicating regions had higher mutation rates (**Fig. 2d**).

For a more detailed analysis, we performed multiple regression. In our analysis, the majority of mutated microsatellites were A/T mononucleotide repeat as previously reported, suggesting that the fragility is primarily determined by the base composition (20) (**Fig. 1b**). Therefore, to find other factors that associate with the mutation rate of microsatellites, we selected 1,000 highly mutated A/T microsatellites of 10-30bp length in the reference genome, and analyzed them for deletions and insertions in the MSS and MSI samples. We considered replication timing, nuclear lamina binding region, G-quadruplexes, and predicted DNA shapes. Nuclear lamina binding regions are known to be associated with genomic fragile sites (26), while G-quadruplexes can cause replication errors (27). The impact of DNA shapes is not well known, but one DNA shape parameter (ORChID2) has been reported to be associated with mutation rate of somatic indels (9). Multiple regression analyses for these factors showed that the length of microsatellite, replication timing, and several DNA shapes were significantly associated with the mutation rate of microsatellites (**Supplementary Table 4**).

The predicted DNA shapes of the flanking sequences were significantly associated with the proportion of mutated samples (**Fig 2e-h and Supplementary Table 4**). Several DNA shape features such as ORChID2 (OH Radical Cleavage Intensity), HelT (Helix Twist), Opening, MGW (Minor Groove Width), Rise, ProT (Propeller Twist), Roll, and Slide, were significantly associated with the prevalence of insertions and deletions in microsatellite regions (**Fig 2e-h, Supplementary Fig. 8, Supplementary Table 4**). The nuclear lamina binding region and G-quadruplexes were not significantly associated (**Supplementary Table 4**). The adjusted *R*^2^ values of the multiple regression analysis were 0.25 in the deletions of MSI, 0.14 in the insertions of MSI, 0.28 in the deletions of MSS and 0.29 in the insertions of MSS (**Supplementary Table 4**).

Microsatellites are highly polymorphic and have also been used as genetic makers for population genetics studies (10). To evaluate the genetic polymorphism, we estimated the heterozygosity of each microsatellite locus in normal tissues. The proportion of mutated samples in cancers and the heterozygosity in normal tissues was significantly correlated (Pearson’s product moment correlation coefficient; *r* = 0.31, *P-value* < 10^-16^) (**Fig 2i**), indicating that genetic variations and somatic mutations are influenced by the same factors.

### Highly-mutated microsatellites

We compared mutability of each microsatellite between the MSI and MSS samples, and selected the top 20 highly-mutated microsatellites with the highest mutation rates (proportion of mutated samples for a microsatellite) (**Supplementary Table 5**). We performed a clustering analysis with these microsatellite markers, and confirmed that they perfectly distinguished the MSI and MSS samples in the CR, UT and ST cancers (**Fig 3a-c**). Although the efficiency of these markers should be evaluated by an independent and larger cohort, we consider that they have a technical advantage over known MSI markers in availability in whole genome sequencing and combinations of the markers can be used as a new marker set.

### Genes with large number of mutated microsatellites

To find genes with high mutation rates (proportion of mutated samples), we tested the total number of indels in microsatellites for each gene across the 21 tumor types. We counted the number of mutated microsatellites and the total number of analyzed microsatellites for each gene, and identified genes with larger numbers of mutated microsatellites compared to others. After adjusting for multiple testing, 1,134 genes had significantly larger numbers of mutated microsatellites for at least one tissue (*q-value* < 0.01) (**Supplementary Table 6**). Of these genes, *ALB,* which is known to be highly expressed in liver, showed the largest number of mutated microsatellite (Fisher’s exact test; *q-value* = 6.5×10^-15^, odds ratio = 65.1) in liver cancer (LI) (**Supplementary Fig. 9**). A previous study suggested that some cell linage-specific highly-expressed genes, including *ALB* in liver, had recurrent short indels (28). This result is consistent with the previous study, and strong DNA damage in cell linage-specific highly-expressed genes would influence mutation rate of microsatellites (**Supplementary Table 6**).

### Recurrently mutated microsatellites in the coding region

To compare recurrently mutated microsatellites in the coding regions between MSS and MSI samples, we calculated the proportion of mutated samples for each microsatellite in coding regions (**Fig 4, Supplementary Table 7**). Microsatellites or repeat sequences in *ACVR2A* and *TGFBR2*, which have been reported to be frequently mutated in MSI tumors (13, 14, 20), were recurrently mutated in 60% and 47% of the MSI samples, respectively. In addition, microsatellites in *ASTE1, KIAA2018, LIN1* and *CDH26* were also mutated in more than 50% of the MSI samples. Mutations in microsatellites in Cosmic Cancer Genes (*MSH6, JAK1, BLM, IL7R,* and *CSF3R*) were identified as MSI specific mutations. Of these, indels in *MSH6,* which is a mismatch repair gene, is likely to cause the MSI phenotype (11). Mutations in *JAK1* in MSI cancers were reported to associate with tumor immune evasion (29). In the MSS samples, microsatellites or repeat sequences in *APC* and *TCF12* were mutated only in MSS samples, suggesting that these mutations cause cancer without genomic instability.

Although many of the recurrently mutated coding microsatellites have been reported by whole genome or exome sequencing studies (**Supplementary Table 7**) (13, 14, 20, 30), our analysis identified new genes with recurrently mutated microsatellites. Of these, the *GINS1* gene encodes a subunit of DNA replication complex (31). *MBD4* has been reported to contribute to tumorigenesis and work as a modifier of MMR-deficient cancer (32). *BLM* is included in the Cosmic Cancer Gene database and has functions in DNA replication and DNA double-strand break repair (33). These results suggest that mutations in microsatellites can work as driver events.

### Mutation in mismatch repair (MMR) and proof-reading genes

We analyzed the association between the proportion of mutated microsatellites, and somatic and germline variants of eight DNA repair genes (*MLH1, MLH3, MSH2, MSH3, MSH6, PMS2, POLE* and *POLD1*). First, we focused on stop gain, splice site, nonsense mutations, and gene-disrupting structural variations in tumor and matched normal samples (**Fig 5a-d**). Two samples in CR and UT had a loss of function germline mutations in the *MSH2* or *PSM2* gene, suggesting that Lynch syndrome could cause cancers in these patients (**Fig 5a and 5c**) (11). The number of samples with somatic SNVs and indels in these genes were significantly enriched in the MSI samples (Fisher’s exact test CR; *P-value* = 2.0×10^-8^, Odds ratio = 10.6, ST; *P-value* = 1.2×10^-4^, Odds ratio = 49.0, UT; *P-value* = 1.4×10^-5^, Odds ratio = 40.6), while those with germline variants and structural variations (SVs) were not significantly enriched. These results suggest that most of the MSI phenotypes in cancer were mainly caused by somatic short indels or somatic SNVs. Germline SVs of *PMS2* were frequently observed in MSS tumors, indicating that *PMS2* could have a lower impact on DNA mismatch repair deficiency or MSI (34).

Most MSI samples had larger numbers of somatic SNVs (**Fig. 1c**), due to functional deficiency of MMR genes. However, 40 MSS samples had larger numbers of somatic SNVs than the average number of SNVs in the MSI samples (151,816.6 SNVs). Of these, 8 had somatic missense mutation in the exonuclease domain of *POLE* (residues 268-471) (35) (**Supplementary Fig. 10, Supplementary Table 8**), suggesting that exonuclease domain mutations of *POLE* were associated with a large number of SNVs in MSS, not MSI.

### Association with somatic mutational signatures in PCAWG

We compared mutational signatures found in single base substitution (SBS), doublet base substitution (DBS), as well as insertions and deletions (ID) between the MSI and MSS samples (**Fig 6a-c, Supplementary Fig. 11, Supplementary Table 9**). The PACWG signature analysis detected 49 SBS, 11 DBS, and 17 ID signatures (36). We compared the fraction of each mutational signature between MSI and MSS samples in CR (Colon/Rectum), ST (Stomach), and UT (Uterus), and found that six SBS signatures (SBS5, SBS15, SBS20, SBS21, SBS26 and SBS44), one ID signature (ID2), and four DBS signatures (DBS3, DBS7, DBS8 and DBS10) were significantly different among the MSI and MSS samples in at least one cancer type (Wilcoxon signed-rank test, *q-value* < 0.05). Except for DBS3 and DBS8, most of these mutational signatures have been reported to be associated with tumors having defective DNA mismatch repair (36). We found DBS3 and DBS8 to be associated with MSI. DBS3 was also associated with the mutations in exonuclease domain of *POLE* in the current study (Wilcoxon signed-rank test, *q-value* < 0.05) (**Supplementary Table 10**), and no etiology has been proposed for DBS8, which was observed in ES (Esophagus adenocarcinoma) and CR (36). In the ID signatures, the fraction of ID2 (A/T deletion) was significantly different between MSI and MSS in CR, ST, and UT, which is consistent with an excess of A/T indels in the microsatellite regions (**Fig.1b**).

### Neo-antigen load from the microsatellite or repeat coding regions

MSI cancers are known to show specific immune reactions such as Crohn-like reaction and diffused infiltration of lymphocytes in pathology (37), and PD-1 inhibiting immune therapy is a highly effective treatment for all types of tumors showing MSI (12). Its specific immune-reaction should be related to neo-antigen burden, and we thus calculated the neo-antigen burdens of MSI and MSS tumors by using somatic mutations and HLA genotypes, taking into account neo-peptides generated from indel or frameshift mutations of the coding microsatellites. The number of predicted neo-antigens in MSI tumors (median, 393) was significantly higher than MSS tumors (median, 11; *P-value* = 5.9×10^-20^) (**Supplementary Fig. 12**), which is consistent with the mutational burden. Interestingly, while 95% of neo-antigens were derived from SNVs in MSS tumors, in MSI tumors, 51% of the predicted neo-antigens were derived from short indels and 5% were derived from indels of the microsatellites.

## Discussion

Due to the clinical importance of MSI phenotypes, exome sequencing and small-scale WGS studies were performed for the MSI samples (13, 14, 20, 30). These studies identified recurrently mutated microsatellites and driver genes mainly located in coding regions, as well as created an algorithm to find MSI with smaller number of microsatellite sets. However, detailed analysis for factors that influence mutation rate and validation of the selected microsatellite marker sets in independent cohorts was limited. Here, we performed an analysis of microsatellite mutations in the largest whole genome sequencing cohort with the largest number of microsatellites to date, so as to characterize microsatellite mutations and MSI tumors at the genome-wide level.

To identify microsatellite regions in the human genome, we used results from three software (Tandem repeat finder, MS detector, and MISA) (16-18). After filtering, we obtained 9,292,677 microsatellites in the genome for analysis. Among the methods for detecting microsatellites, the MISA software identified the largest number of microsatellites (17). Although most of them were short and may not be considered as microsatellites by the other two methods, they contained highly-mutated repeat regions. It has been reported that the rate of mutation of longer microsatellites is higher than that of shorter microsatellites (38). However, current short-reads sequencing technologies are unable to analyze longer microsatellites. Indeed, we could not obtain sufficient number of reads for Bethesda markers (**Supplementary Fig. 7**). We therefore decided to prioritize shorter repeats as the main targets for the current WGS study. Alternatively, in this study, we detected highly-mutated short microsatellites (**Supplementary Table 5**) and they could be useful for clinical diagnosis of MSI with current short read technologies.

The analysis of WGS and the validation study in an independent cohort found 20 novel microsatellite markers (**Fig. 3**), which can be used to predict tumors with high mutational burden. However, the mutation rate of microsatellite was highly correlated with the total number of indels, but not strongly correlated with the total number of SNVs (**Fig. 1cfg**), indicating that samples with larger numbers of SNVs can be found in MSS samples. As the analysis of neo-antigens showed that SNVs can also produce a larger number of neo-antigens (**Supplementary Fig. 11**), identification of these samples is also important for diagnosis. The analysis of MSS samples with a larger number of SNVs showed that mutations in the exonuclease domain of *POLE* can partly explain the high mutation rate of SNVs, instead of indels and microsatellites (**Supplementary Fig. 10**). Therefore, analysis of mutations in the *POLE* gene in MSS samples can identify more tumors with high mutational burdens (39).

Our WGS analysis found high rates of short deletions in the microsatellites of both MSI and MSS samples (**Fig. 1de**). The excess of deletion events was also observed in previous studies (20). Therefore, the excess of short deletions should not be due to a bias of our mutation calling method, and can be considered as a common feature of cancers. A microsatellite mutation model suggests that deletions are generated by a misalignment loop in the template strand, and insertions subsequently generated in the nascent strand (10). During the DNA replication of cancer cells, template strands could exist as single strands for a longer period compared to nascent strands, resulting in a higher chance to generate misaligned loop structures, which would induce larger numbers of deletions.

The analysis of replication timing showed a different pattern between deletions in MSI and others (**Fig. 2a-d, Supplementary Table 4**). As observed in the SNVs, replication timing and mutation rate were positively correlated with the insertions and deletions of MSS samples, as well as insertions of MSI samples (**Fig. 2a-d**). It is suggested that early replicating regions are more accessible for DNA repair machineries, resulting in more chances for repair (8). However, deletions in microsatellites were enriched in the early replicating regions of MSI samples **(Fig. 2d)**. A recent exome sequencing study also reported the inverse correlation between the microsatellite indels and replication timing in MSI tumors (20). Since MSI tumors should have defects in their DNA mismatch repair machinery, this result would reflect the pattern of mutation without DNA mismatch repair. In early replication, template strands may exist as single strands for a longer period, facilitating the occurrence of deletions.

In addition to replication timing, DNA shape parameters were also associated with the mutation rate of microsatellites. Microsatellites with lower HelT (Helix twist), higher ProT (Propeller twist), higher Roll, and higher Slide had higher insertion rates (**Fig. 2e-h, Supplementary Fig. 8**). Microsatellites with higher HelT, higher Opening, higher ProT, higher Roll, and lower Slide had higher deletion rates (**Fig. 2e-h, Supplementary Fig. 8**). Since DNA shapes were associated in both the MSI and the MSS tumors, they would affect the fragility of DNA strand, and mainly influence the mutation generation instead of the repair process.

In the present study, we considered that the mutation rate of microsatellites is mainly influenced by the following; fragility of DNA sequences (length and unit type of microsatellite, and DNA shape) (**Fig. 1b**, **Fig. 2e-h**), activity of DNA repair machinery (mutations or activities in mismatch repair machinery genes) (**Fig. 5)**, DNA damage against cell linage-specific highly-expressed genes (**Supplementary Fig. 10, Supplementary Table 6**), and accessibility of DNA repair machinery (DNA replication timing) (**Fig. 2a-d**). Furthermore, since the mutation rate in cancer was correlated with the heterozygosity of germline variations, these factors would also affect the mutation rate in germline variations (**Fig. 2i**).

The current study analyzed somatic indels in microsatellite regions in the largest WGS cohort to date. We found a microsatellite marker set to detect MSI, factors that influence the mutation rate of microsatellite, genes with recurrently mutated microsatellites, and the influence of somatic mutations in MMR and proof-reading genes have on MSI. Our analysis provides a mutational landscape of microsatellites in cancer samples for future clinical applications.

## Acknowledgements

The super-computing resource “SHIROKANE” was provided by Human Genome Center, The University of Tokyo (http://sc.hgc.jp/shirokane.html). This work was supported partially by Grand-in-aid for RIKEN CGM and IMS, Grant-in-Aid for Scientific Research on Innovative Areas from JSPS grants (25134717, 25670375, 23114001, 15H04814), Project for Cancer Research and Therapeutic Evolution (P-CREATE) (Grant Number 16cm0106519h0001, to H, N.). and Platform Program for Promotion of Genome Medicine (Grant Number 18km0405207h0003, to A.F.) in the Japan Agency for Medical Research and Development (AMED).

## URLs

Software for somatic mutation; MIMcall (https://github.com/afujimoto/MIMcall)

Software for germline variation; MIVcall (https://github.com/afujimoto/MIVcall)

DNA shape parameters; http://rohsdb.cmb.usc.edu/GBshape/

Lamina binding region; https://www.nature.com/article-assets/npg/nature/journal/v453/n7197/extref/nature06947-s2.txt Software for estimating location of G-quadruplex;

https://github.com/dariober/bioinformatics-cafe/blob/master/fastaRegexFinder.py

Replication timing; https://genome.ucsc.edu/cgi-bin/hgFileUi?db=hg19&g=wgEncodeUwRepliSeq

Cosmic cancer genes; https://cancer.sanger.ac.uk/cosmic

R; https://www.r-project.org

## Author Contributions

Study design: A.F., S.G.R., M.R.S., and H.N. Data analysis: A.F., M.F., H.T., Y.S., S.M. S.I. and H.N. Molecular analysis: K.M., A.O-S., K.N. and H.N. S. Manuscript writing: A.F., M.F. and H.N.

## Materials and Methods

### Samples and data

Whole genome sequencing data was obtained by the International Cancer Genome Consortium (ICGC) pan-cancer project (15). The list of analyzed samples is shown in the **Supplementary Table 1 and 3**. Datasets of somatic point mutations, short indels, structural variants (SVs), and copy number alterations were generated as part of the Pan-Cancer Analysis of Whole Genomes (PCAWG) project (21, 22). Overall, 2,834 samples with whole genome data are represented in the PCAWG datasets, spanning a range of cancer types (bladder, sarcoma, breast, liver-biliary, cervix, leukemia, colorectal, lymphoma, prostate, esophagus, stomach, central nervous system, head/neck, kidney, lung, melanoma, ovary, pancreas, thyroid, and uterus). The consensus somatic SNVs and short indels in PCAWG samples were determined using different algorithms; calls made by at least two algorithms were used in downstream analyses (22).

### Definition of microsatellite region for the analysis

We determined microsatellite regions using MsDetector, Tandem Repeat Finder and MISA software (16-18). Microsatellite regions defined by the Tandem Repeat Finder were obtained from the UCSC database (16). Identification of microsatellite with MISA was done by (unit size) = 1 to 5, (minimum number of repeats) = 5 and (max difference between 2 microsatellites) = 10. Since these three methods used different algorithms to define microsatellites, we first defined the repeat unit of each microsatellite. We divided each region by different lengths (1-6bps), and calculated the entropy of the character string. The length with the lowest entropy was selected as the unit length of each microsatellite region. For the analysis of the microsatellite, we filtered microsatellite regions according to the following criteria; (1) the proportion of the most frequent unit ≥ 0.8, (2) distance between closest neighboring microsatellite ≥ 30bp, (3) if the microsatellite regions were detected by 2 or more methods, we selected the longest one and discarded others, and (4) upstream and downstream flaking sequences (100bp) of each microsatellite were mapped against human reference genome (GRCh37) by blat software (40) with the options of -stepSize=5 and -repMatch=2253, and microsatellites that had ≥ 90bp of flanking sequences mapped to different positions were removed. As a result of this selection procedure, 8,817,054 autosomal microsatellites remained and were used for the subsequent analyses (**Supplementary Fig. 1**).

### Error rate estimation of each repeat unit

To identify somatic indels in the microsatellite regions, we first estimated the error rate of the different types of repeat units. The type of microsatellites was defined by length of microsatellite region in the reference genome and repeat unit. Microsatellites were categorized by length (6-9, 10-19, 20-29, 30-39, and 40-100bp) and repeat unit and error rates were estimated for each category (see **Supplementary Table 2**). For this purpose, we used data from chromosome X of 32 male normal samples, because chrX is homozygotic and error rates can be estimated without the influence of heterozygous polymorphisms (19, 20). The estimated error rates are shown in **Supplementary Fig. 2**.

### Identification of change of repeat unit from whole genome sequence

Microsatellites are repeat sequences and mapping errors can influence the accuracy of detection. To remove possible mapping errors, we removed improper pairs and reads with low mapping quality (< 30), as well as reads with large (> 550bp) or small (< 100bp) read pair distance.

We counted the number of repeat units in each microsatellite region. We then determined the genotype of the matched normal tissues and detected somatic indels by comparing the genotype of the normal and cancer samples. To distinguish the mutation or variation from sequencing errors, we incorporated the binomial distribution with the estimated error rates (**Supplementary Fig. 2**) and calculated a likelihood for each variant candidate.

For normal samples, we calculated the likelihood for the second most frequent number of repeat (**Supplementary Fig. 2**).

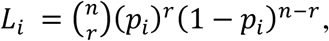

where *n* is the total number of reads that cover the microsatellite, *r* is the number of reads containing *i*th repeat, and *p*_*i*_ is the estimated error rate of the *i*th repeat. If the likelihood is lower than a threshold value, the genotype was assumed to be heterozygous for the major repeat and second major repeat. We next calculated the likelihood for the number of repeats in the cancer. If the likelihood of the non-germline repeat was lower than a threshold value, we defined the repeat as a somatic indel candidate.

To find the appropriate likelihood threshold values, we applied this algorithm on data from other male chrX. Based on the estimated false positive and negative rates, we set −1 and −8 for germline genotyping and somatic mutation calling (**Supplementary Fig. 3**) respectively. Based on the comparison, we set *L*=-8 for tumor, and *L*=-1 for matched normal samples. We selected microsatellites that were covered by ≥ 15 reads in both the cancer and matched normal samples. Additionally, we selected somatic indels with variant allele frequencies in cancer ≥ 0.15 and number of support reads in cancer ≥ 2 and ≤ 1 in normal samples.

### Estimation of false discovery rate

We randomly selected 29 somatic MS mutations detected in liver cancer samples RK001, RK249 and RK308 and performed validation with a previously reported method (41). Amplicons were analyzed using the ABI PRISM ® 3100 Genetic Analyzer (Applied Biosystems), and GeneMapper software (Applied Biosystems). Validation for the selected microsatellites was also performed using the Sanger sequencing method (**Supplementary Fig. 5**).

### Selection of highly-mutated MSs

To select microsatellites, we compared the number of mutated samples in microsatellite instability (MSI) and microsatellite stable (MSS) samples for each microsatellite using Fisher’s exact test. Nine microsatellites with odds ratio ((number of mutated samples in the MSI)/ (number of un-mutated samples in the MSI)/ (number of mutated samples in the MSS)/ (number of un-mutated samples in the MSS)) ≥ 500 and proportion of the mutated samples in the MSI samples ≥ 0.8 were selected and genotyped in the additional samples. Three microsatellites were selected from highly-mutated microsatellites in the MSI samples. Ten microsatellites were selected from recurrently mutated coding microsatellites.

### Threshold determination for MSI

Since the mutation status of conventional MSI markers could not be obtained (see **Supplementary Fig. 7**), we needed to determine the threshold value to select MSI. We first excluded colorectal, stomach and uterus cancers (n=186) from all samples (n=2717). We assumed that the other cancers contained negligible number of MSI samples. We then calculated the average and standard deviation of the mutation rates. We also assumed that the distribution of the mutation rates follows a normal distribution with the obtained average and standard deviation. The 99.99th percentile of the normal distribution was 0.0254. Therefore, we adapted 0.03 (slightly conservative value from the 99.99th percentile) for the criteria for MSI in genome-wide level, and 31 samples were defined as MSI in this study. No colorectal, stomach or uterus cancers were used to determine the threshold value, however, the value still gave a reasonable grouping for colorectal, stomach and uterus cancers (please see the **Fig. 1 (b)**).

### Comparison of mutational signatures between the MSS and MSI

Mutational signatures and their proportions were obtained from the result of the PCAWAG signature working group (36). The proportion of each signature was compared between MSI and MSS samples in CR, UT, and ST with the Wilcoxon signed-rank test. Multiple testing adjustment was done using Benjamini and Hochberg’s FDR method (42).

### Identification of genes with recurrently mutated microsatellites

To identify highly-mutated genes, we compared the mutation rate of microsatellites in each gene. We compared the total number of analyzed microsatellites and total number of mutated microsatellites in introns and exons in each gene. We also counted the total number of analyzed microsatellites and total number of mutated microsatellites in the entire genome in MSS samples for each cancer type; (total number of mutated microsatellites in gene *i* in all MSS samples in cancer *j*)/(total number of un-mutated microsatellite in gene *i* in all MSS samples in cancer *j*) and (total number of mutated microsatellite in entire genome in all MSS samples in cancer *j*)/(total number of un-mutated microsatellite in entire genome *i* in all MSS samples in cancer *j*) and these were compared with Fisher’s exact test. Multiple testing adjustment was done using Benjamini and Hochberg’s FDR method (42), and from this analysis, we could obtain genes with larger numbers of mutated microsatellites compared to the entire genome.

### Analysis of epigenetic factors on mutability of microsatellite

To find the factors that influence the mutability of microsatellites, we considered replication timing, nuclear lamina binding region, G-quadruplexes, and predicted DNA shapes (See URLs section). For the replication timing, we downloaded data of HepG2, K562, MCF-7, SK-N-SH, and GM12878 cells. We averaged the replication timing within 1Mbp bins for each cell line and bins with standard deviation ≤ 15 were used for the analysis. Presence or absence of nuclear lamina binding region and G-quadruplexes within ±1000bp from the start and end of microsatellites were examined. The predicted DNA shapes (Buckle, HelT, minor groove width (MGW), ORChID2, Opening, ProT, Rise, Roll, Shear, Shift, Slide, Stagger, Stretch and Tilt) of ±5bp from the start and end of each microsatellite were used for the analysis. We performed a multiple regression analysis of the parameters with lm function of R software, and parameter selection was done with step() function.

### Prediction of neo-antigens

HLA genotyping from WGS data were generated as part of the PCAWG project. Somatic point mutations, non-MS indels detected by the PCAWG project, and MS indels detected by our method were combined and annotated using ANNOVAR. Mutant peptides of length 8–11 residues were assessed for their binding affinity (IC_50_) to the HLA class I of matched patients using NetMHCpan-3.0 (43). Mutant peptides of IC_50_< 50 nM were predicted as neo-antigens.

