## Supplementary material for "Comprehensive Analysis of Indels in Whole-genome Microsatellite Regions and Microsatellite Instability across 21 Cancer Types": Supplymentary_information

**Supplementary Figure 1. Definition of microsatellite (MS) in the human genome.**
**Supplementary Figure 2. Error rate estimation with hemizygotic chrX data.**
**Supplementary Figure 3. Evaluation of threshold.**
**Supplementary Figure 4. Comparison with PCAWG callers.**
**Supplementary Figure 5. Validation of somatic indels in microsatellites.**
**Supplementary Figure 6. Proportion of mutated microsatellites among different categories.**
**Supplementary Figure 7. Mutation rate of Bethesda markers.**
**Supplementary Figure 8. DNA shape and mutability of microsatellites.**
**Supplementary Figure 9. Analysis of microsatellite mutation rate per gene.**
**Supplementary Figure 10. Mutation rate and *POLE*-mutated samples.**
**Supplementary Figure 11. Mutation rates of microsatellites and PCAWG mutational signatures.**
**Supplementary Figure 12. Prediction of neo-antigens in MSI/MSS tumors.**

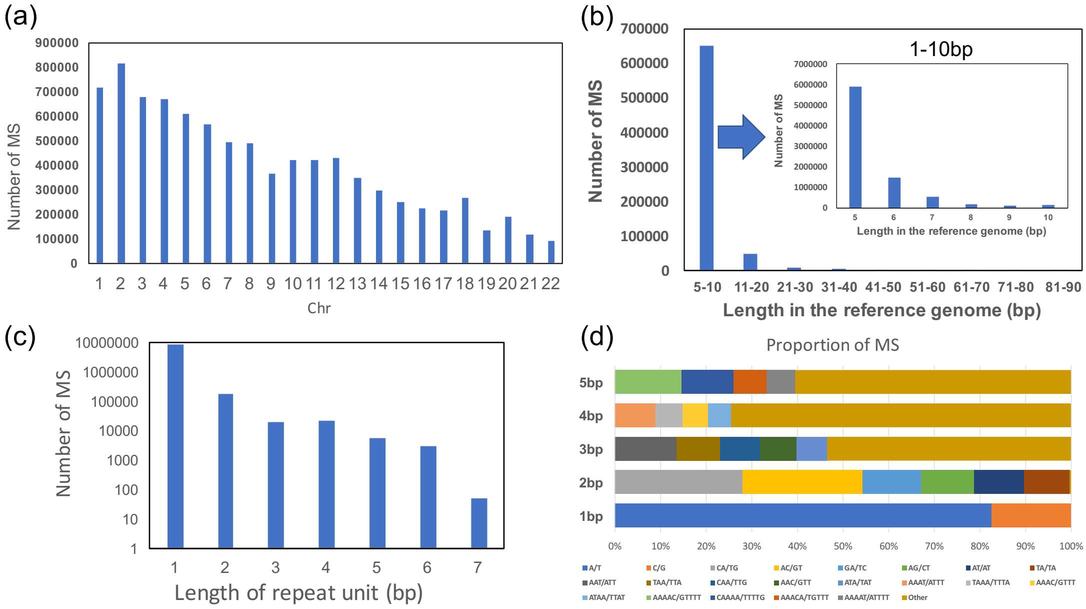


**Supplementary Figure 1. Definition of microsatellite (MS) in the human genome.** (a) Number of microsatellites in each chromosome. (b) Length distribution of microsatellites. (c) Distribution of unit length. (d) Proportion of MS type.

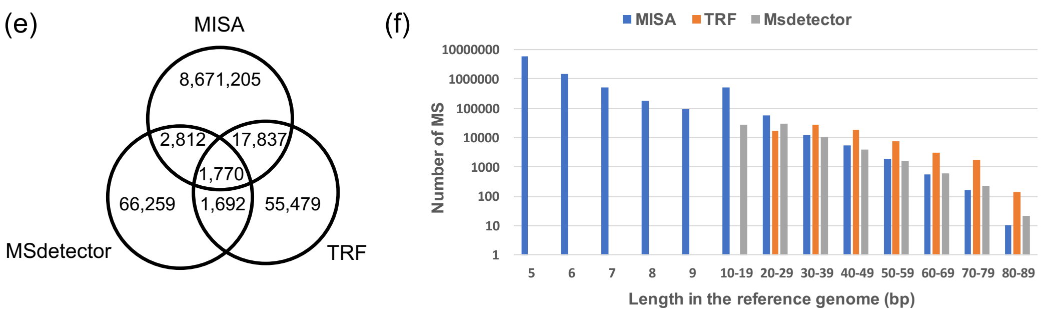


**Supplementary Figure 1. Definition of microsatellite (MS) in the human genome.** (continued) (e) Number of MS regions by different methods. (f) Length distribution of MS in different methods.

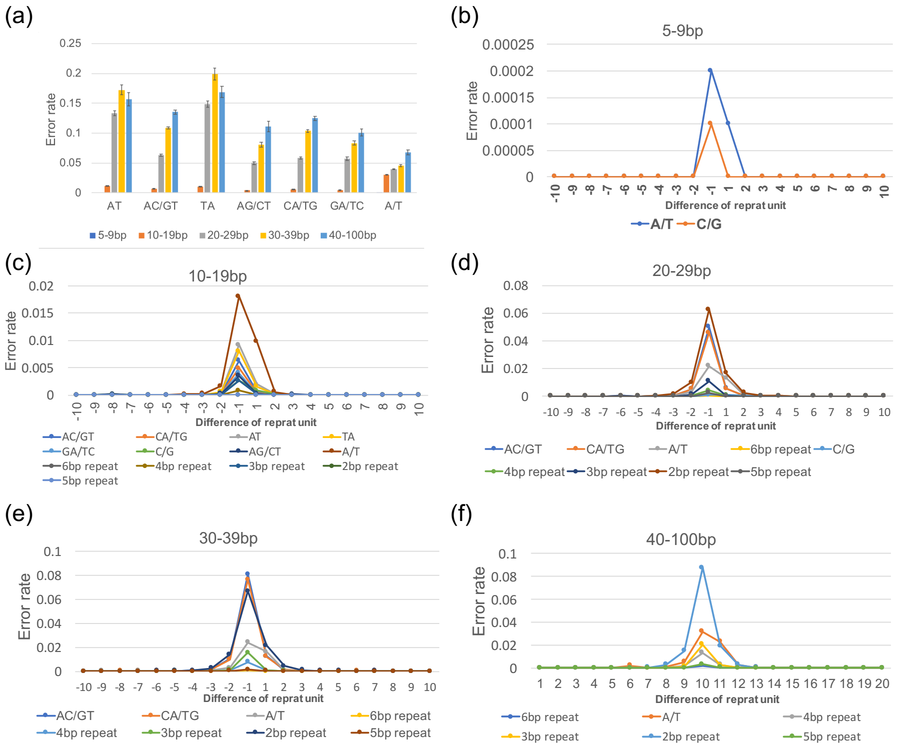


**Supplementary Figure 2. Error rate estimation with hemizygotic chrX data.** (a) Error rate/reads for different microsatellite length and unit type. (b) Error rate of microsatellites 5-9bp in length. (c) Error rate of microsatellites 10-19bp in length. (d) Error rate of microsatellites 20-29bp in length. (e) Error rate of microsatellites 30-39bp in length. (e) Error rate of microsatellites 40-100bp in length.

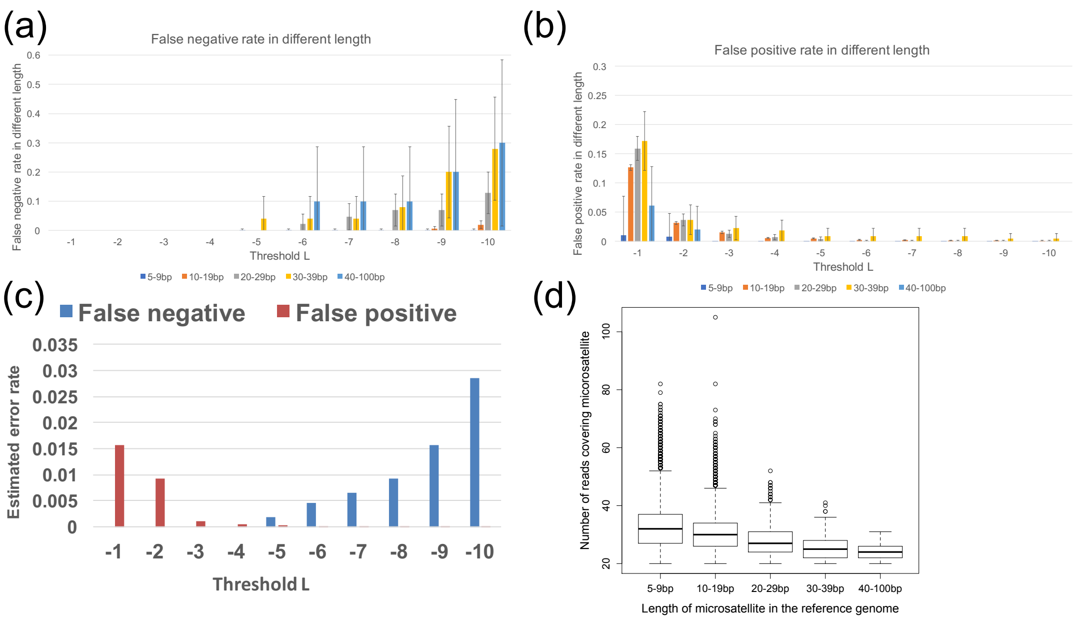


**Supplementary Figure 3. Evaluation of threshold.** (a) Threshold for microsatellite mutation calling and false negative rate. Higher *L* value showed higher false negative rate. (b) Threshold for microsatellite mutation calling and false positive rate. Higher *L* value showed lower false positive rate. (c) Threshold for microsatellite mutation calling and false positive and negative rates. Trade-off effect of *L* values are shown. (d) Number of reads covering microsatellites.

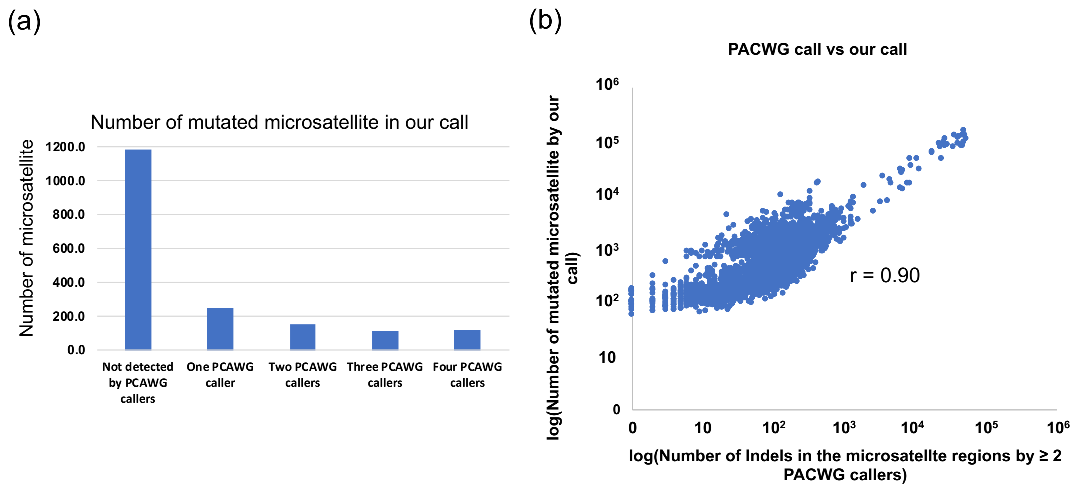


**Supplementary Figure 4. Comparison with PCAWG callers.** (a) Number of identified mutated microsatellites with MIMcall and PCAWG callers. (b) Correlation between our call and consensus with PCAWG callers ( ≥ 2 callers).

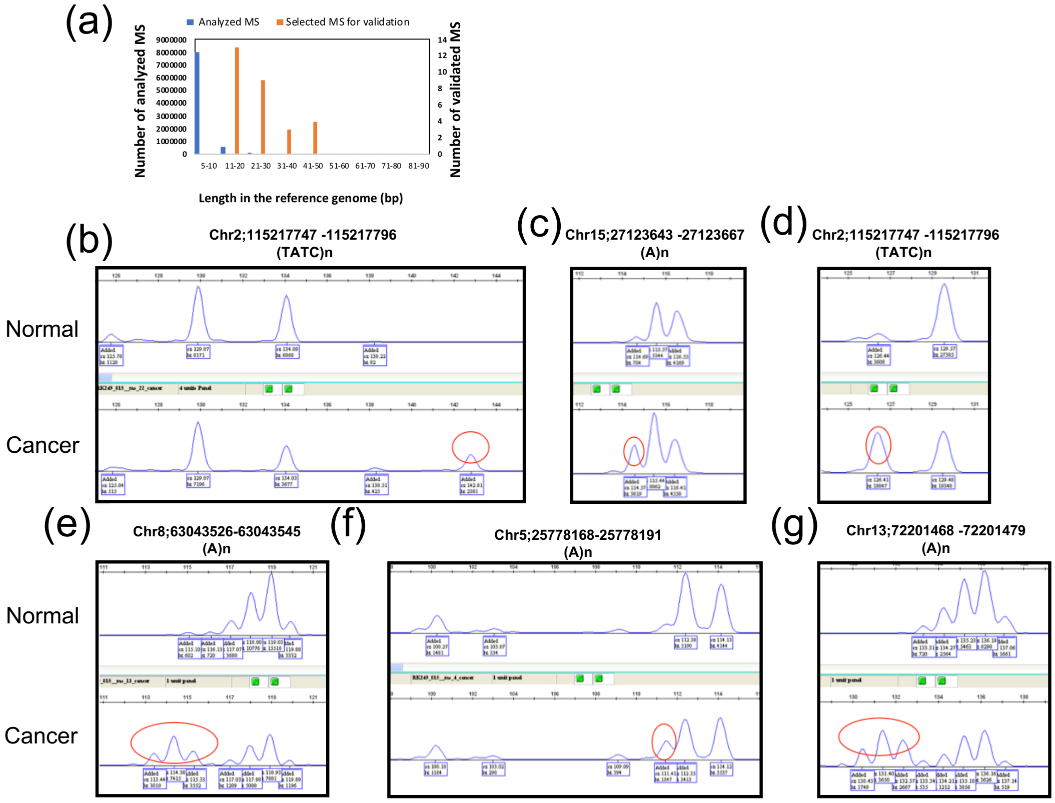


**Supplementary Figure 5. Validation of somatic indels in microsatellites.** (a) Length distribution of all microsatellite (MS) and MS selected for validation. (b-g) Examples of the electrophoresis are shown.

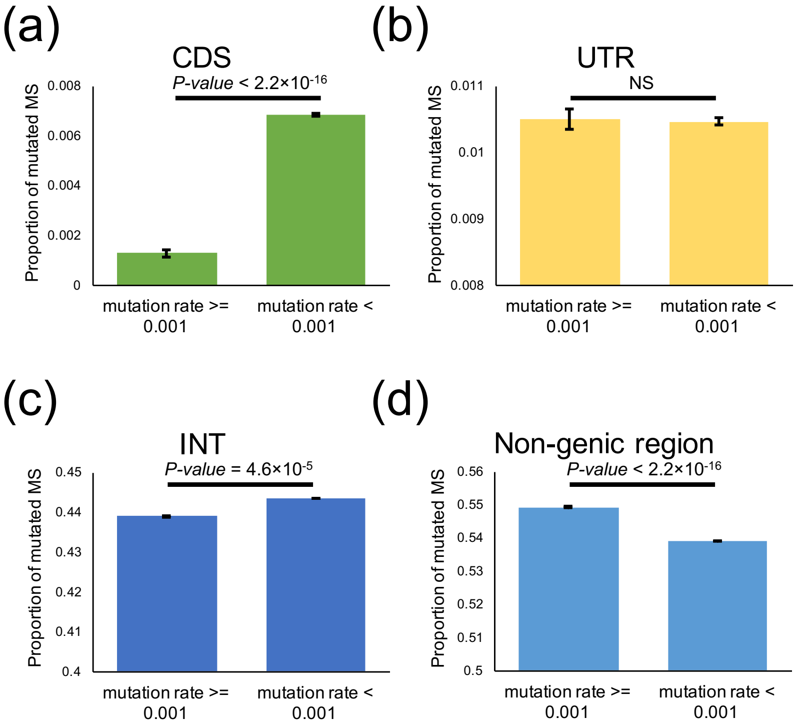


**Supplementary Figure 6. Proportion of mutated microsatellites among different categories.** (a) Coding region. (b) UTR region. (c) Intronic regions. (d) Non-genic region. Numbers of microsatellites with mutation rate ≥ 0.001 and those with < 0.001 were compared for each category with chi-squared test.

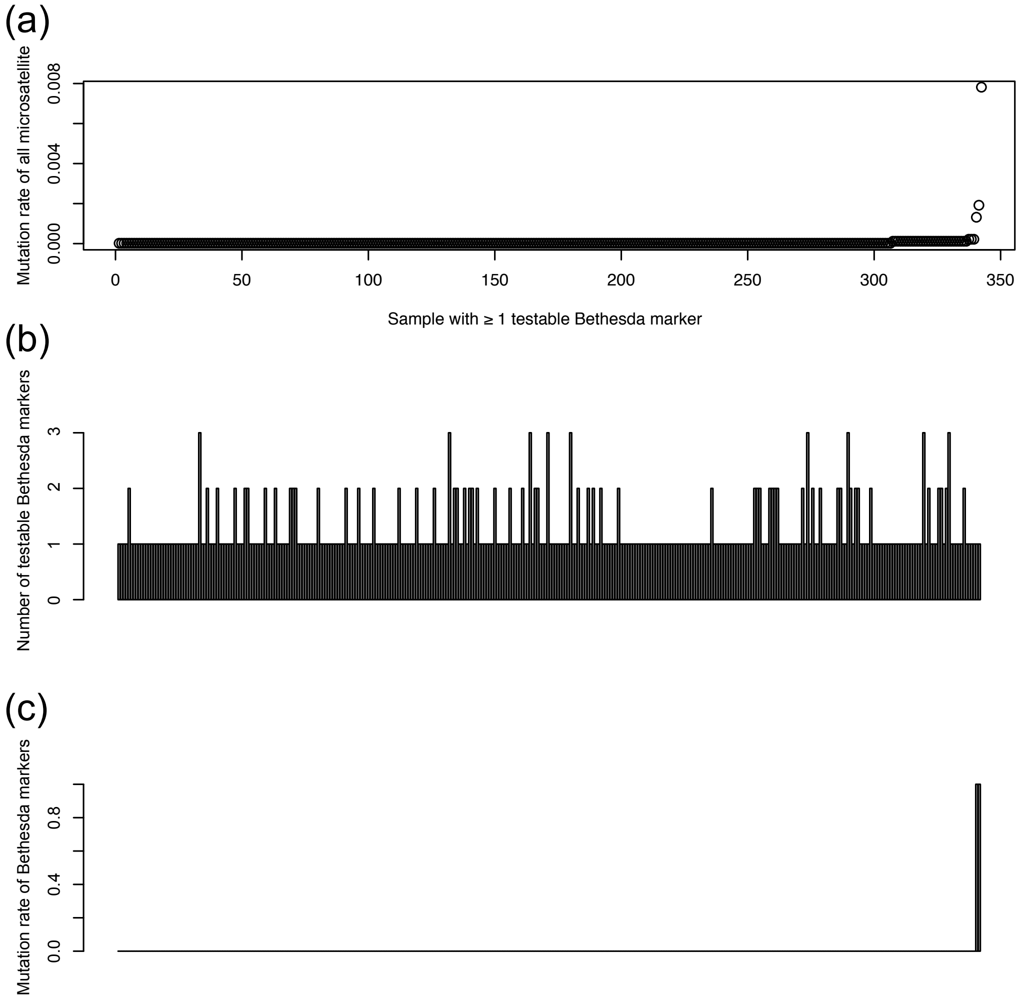


**Supplementary Figure 7. Mutation rate of Bethesda markers.** (a) Mutation rate of all microsatellites of samples with ≥ 1 testable Bethesda markers (number of reads ≥ 15 in cancer and matched normal samples). Only 342 samples had ≥ 1 testable Bethesda markers. (b) Number of testable Bethesda markers in each sample. (c) Mutation rate of Bethesda markers in each sample. Pattern of mutation rate of all microsatellites and Bethesda markers were consistent.

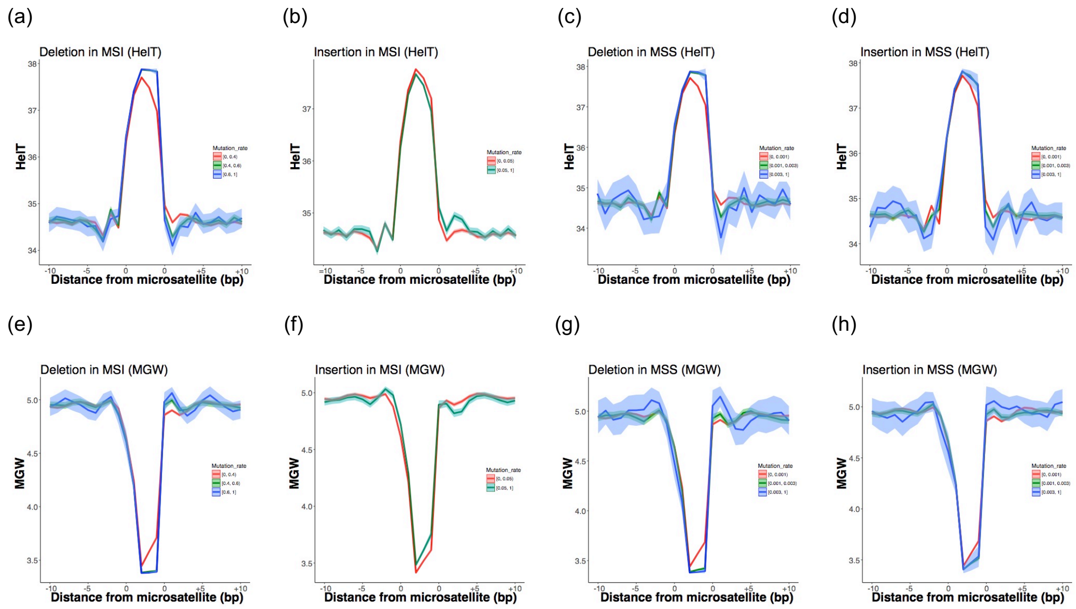


**Supplementary Figure 8. DNA shape and mutability of microsatellites.** Distance from microsatellites and HelT for microsatellites with different mutability; (a) Deletion in MSI, (b) Insertion in MSI, (c) Deletion in MSS and (d) Insertion in MSS. Distance from microsatellites and minor groove width (MGW) for microsatellites with different mutability; (e) Deletion in MSI, (f) Insertion in MSI, (g) Deletion in MSS and (h) Insertion in MSS. Average and 95% confidence intervals are shown.

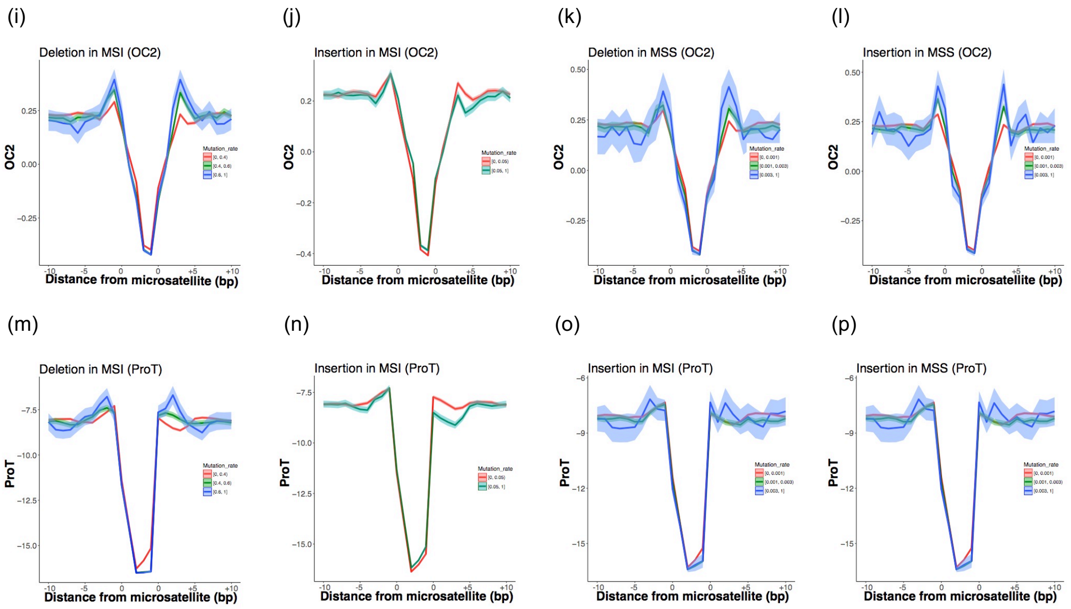


**Supplementary Figure 8. DNA shape and mutability of microsatellites.** (continued) Distance from microsatellites and OC2 (ORChID2) for microsatellites with different mutability; (i) Deletion in MSI, (j) Insertion in MSI, (k) Deletion in MSS and (l) Insertion in MSS. Distance from microsatellites and ProT for microsatellites with different mutability; (m) Deletion in MSI, (n) Insertion in MSI, (o) Deletion in MSS and (p) Insertion in MSS. Average and 95% confidence intervals are shown.

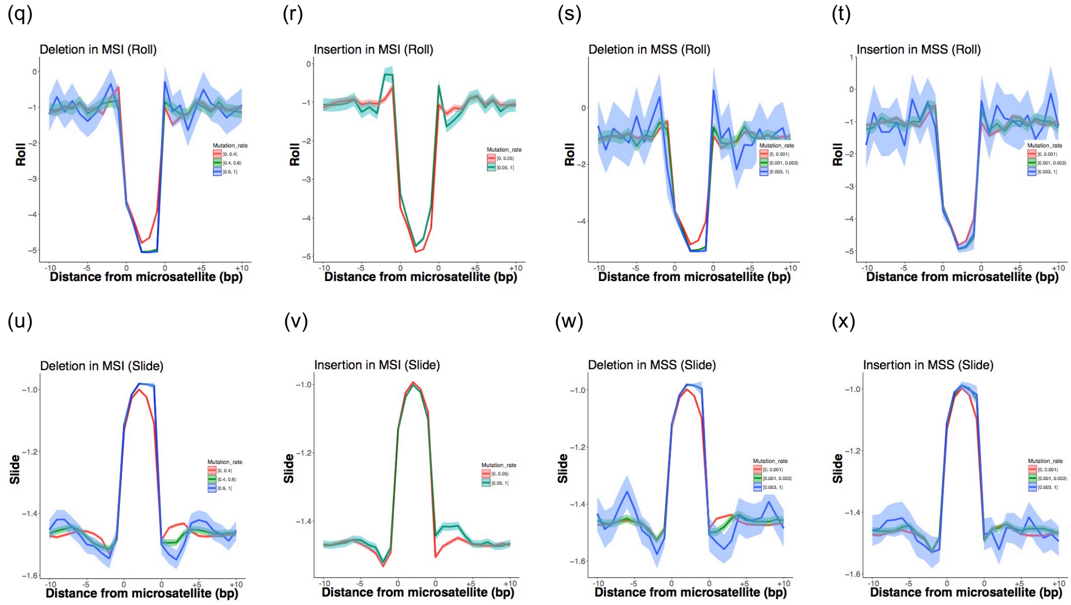


**Supplementary Figure 8. DNA shape and mutability of microsatellites.** (continued) Distance from microsatellites and Roll for microsatellites with different mutability; (q) Deletion in MSI, (r) Insertion in MSI, (s) Deletion in MSS and (t) Insertion in MSS. Distance from microsatellites and Slide for microsatellites with different mutability; (u) Deletion in MSI, (v) Insertion in MSI, (w) Deletion in MSS and (x) Insertion in MSS. Average and 95% confidence intervals are shown.

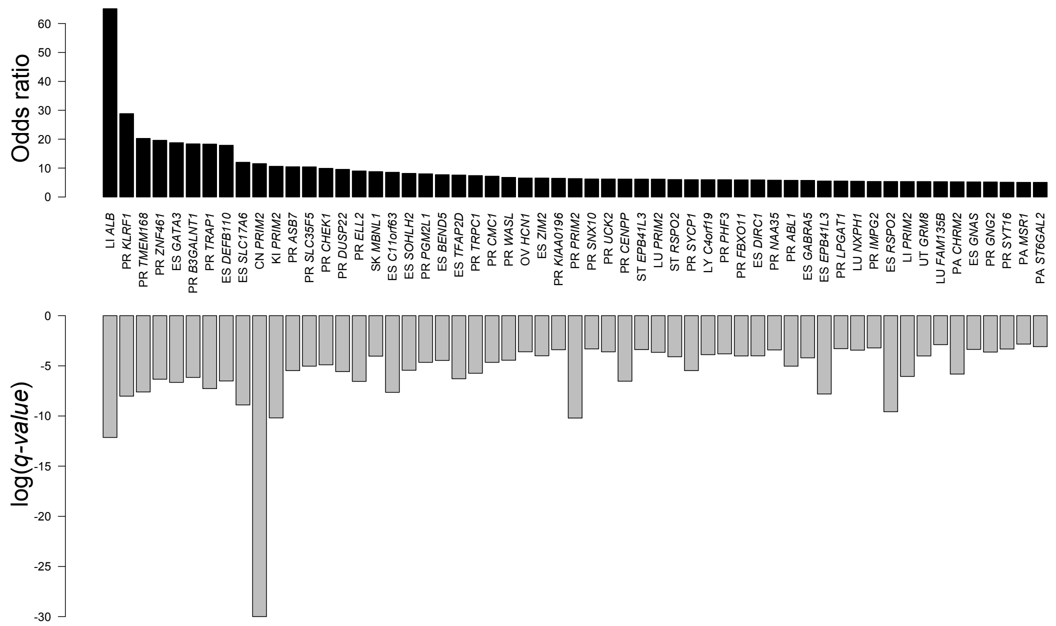


**Supplementary Figure 9. Analysis of microsatellite mutation rate per gene.** Total number of mutated microsatellites was calculated and compared with those of other genes. Genes with total number of mutated microsatellites ≥ 10 were used for analysis. Genes with odds ratio ≥ 5 are shown in this figure. Cancer types and gene names are shown. Upper panel; Odds ratio = (total number of mutated microsatellites in a gene)/(total number of un-mutated microsatellites in a gene)/ (total number of mutated microsatellites in all other genes)/(total number of un-mutated microsatellites in all other genes). Lower panel; log(*q-value*) of the number of mutated microsatellites.

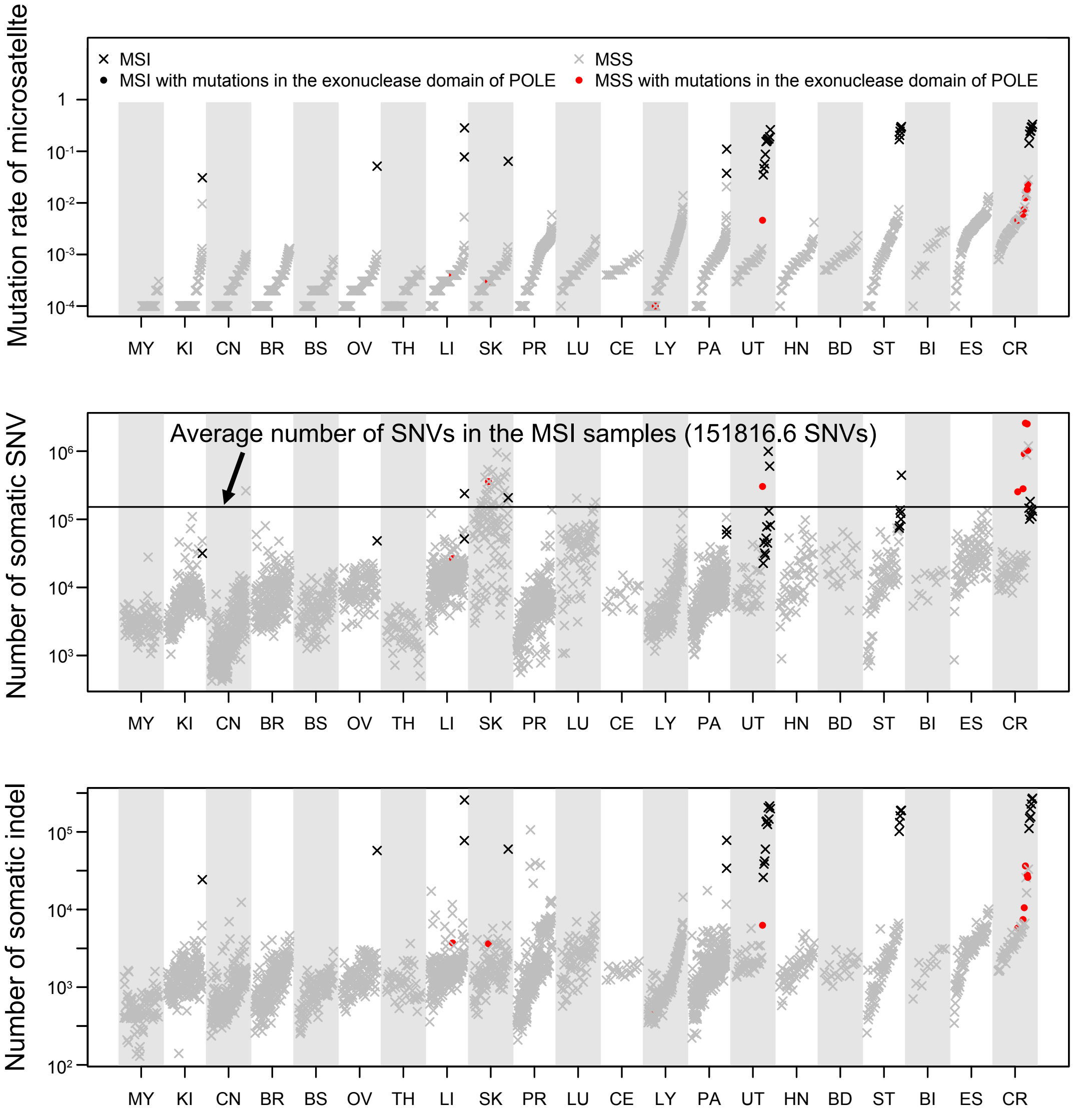


**Supplementary Figure 10. Mutation rate and *POLE*-mutated samples.** Mutation rates of MSI, MSS and samples with mutations in the exonuclease domain of *POLE* are shown. Upper panel; Mutation rate of microsatellites, Middle panel; Number of somatic SNV, and Lower panel; Number of somatic indels.

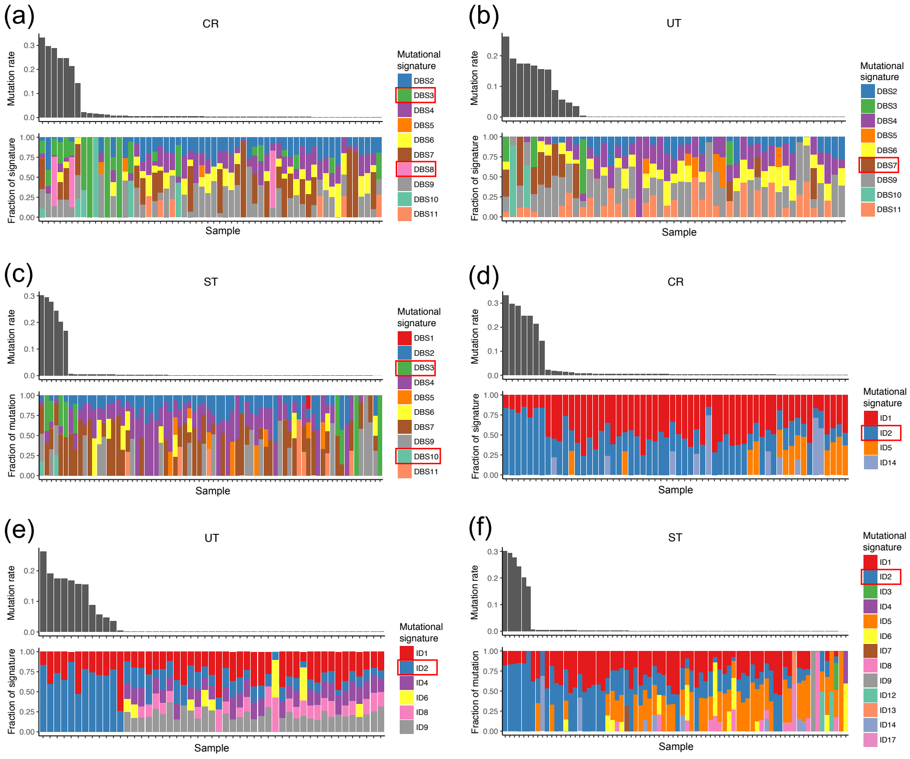


**Supplementary Figure 11. Mutation rates of microsatellites and PCAWG mutational signatures.** Mutation rate of microsatellites and fraction of mutational signatures are shown. Mutational signatures that differed significantly between the MSI and MSS samples were squares in figure legends. (a) DBS (double base substitution) signature of CR, (b) DBS signature of UT, (c) DBS signature of ST, (d) ID (indel) signature of CR, (e) ID signature of UT, (f) ID signature of ST.

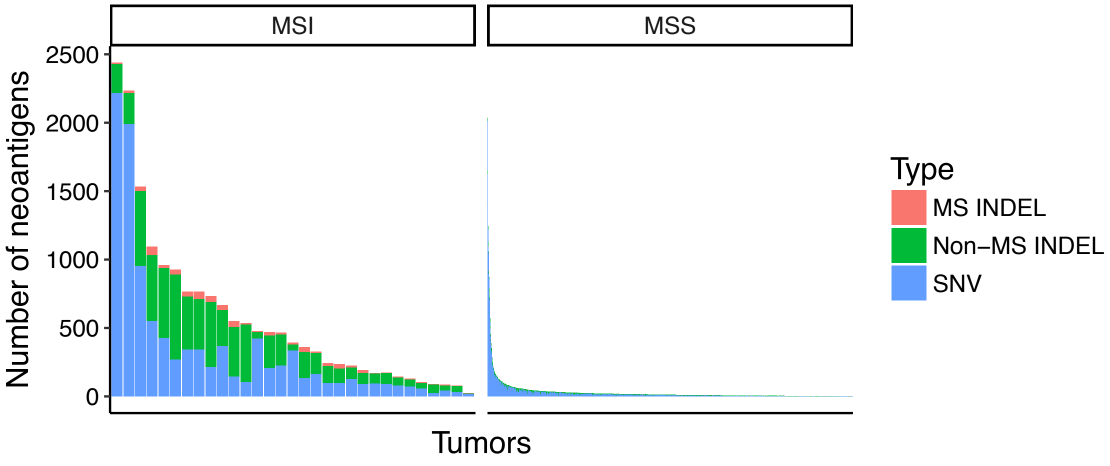


**Supplementary Figure 12. Prediction of neo-antigens in MSI/MSS tumors.** The numbers of neo-antigens were compared between 31 MSI samples and 2,531 MSS samples.
